# Salmonberry Transcriptome Reveals Phylogeny and Novel Badnavirus Species

**DOI:** 10.64898/2026.06.03.727011

**Authors:** Alex L. Salley, Nithish Narasimman, Adithi Raghavan, Ananya Rajagopalan, Siddharth Chandrasekar, Harlow M. Graves, Amir Zur, Maxwell Sherman, Eric Marnadi, Jae Geller, Thelma F. Madzima, Mohua Bose, Manoj P. Samanta

## Abstract

The Rosaceae family comprises thousands of species across over 100 genera, including Salmonberry (*Rubus spectabilis*), a Pacific Northwest native within the diverse *Rubus* genus. Its berries and leaves are used for food and medicinal purposes, and ecologically it functions as a pioneer species that supports biodiversity and limits erosion. Although many *Rubus* genomes were sequenced and analyzed, salmonberry remains undercharacterized: despite a recently sequenced genome, no publicly available annotation or gene expression analysis currently exists. Here, we used RNA sequencing to characterize the salmonberry leaf transcriptome and examine its phylogenetic relationship within *Rubus*. The assembled 63,285 unique transcripts included 1,389 high-confidence lncRNA transcripts expressed in salmonberry leaves, 218 of which are conserved across *Rubus*. Phylogenetic analysis indicates that salmonberry is closely related to *Rubus arcticus*. In addition, we detected a novel species of virus associated with salmonberry. These findings provide foundational genomic resources for *R. spectabilis* and offer new insights into its evolutionary relationships and endogenous viral integrations.

## Introduction

Salmonberry (*Rubus spectabilis*) is an edible bramble native to the Pacific Northwest, ranging from southern Alaska to northern California, where it thrives in moist coastal environments. Salmonberry produces orange to deep red aggregate fruits and grows as a shrub with prickly stems. A member of the *Rubus* genus (raspberry, blackberry) within the Rosaceae, it is closely related to economically important crops such as apples, pears, cherries, peaches, and plums. Beyond its botanical significance, salmonberry holds rich cultural and ecological value. Indigenous Pacific Northwest communities and local populations traditionally consumed the berries, brewed teas from the leaves, and applied leaf preparations for medicinal purposes (GRuB: Garden Raised Bounty; Joseph 2017). Wildlife species rely on salmonberry as a food source and forage, making it an integral component of regional ecosystems.

As a diploid species, *Rubus spectabilis* provides a tractable model for genomic and evolutionary studies within the taxonomically complex genus *Rubus*, which comprises more than 700 species distributed across multiple subgenera. Recent genomic studies have begun to clarify evolutionary relationships within the group (Alice and Campbell 1999; Huang et al. 2023; Gao et al. 2023). While DNA-based analyses reveal genetic structure and ancestry, transcriptomic approaches provide complementary insights into actively expressed genes associated with development, physiology, environmental response, and secondary metabolism. RNA sequencing (RNA-Seq) has been widely applied to related *Rubus* species, including raspberry and blackberry fruits and leaves, identifying genes involved in fruit development, ripening, stress responses, viral interactions, and metabolic regulation (Garcia-Seco et al. 2015; Yang et al. 2021; Chen et al. 2021; Hytönen et al. 2018; Gutierrez et al. 2017; He et al. 2024; Zhang et al. 2024). However, despite the ecological importance of *Rubus spectabilis* and its close relationship to cultivated relatives, its transcriptome has not previously been characterized.

Here, we present a transcriptomic analysis of salmonberry leaves using next-generation sequencing. We annotated an unpublished salmonberry genome obtained from the Canadian BioGenome Project and used it as the reference for transcriptome annotation. We identified expressed genes, examined phylogenetic relationships within the genus *Rubus*, and detected novel viral sequences associated with this species. Together, these results provide foundational genomic resources for *Rubus spectabilis* and contribute to a broader understanding of gene expression and evolutionary relationships within the genus.

## Materials and Methods

### Plant Material and RNA Extraction

On Oct 21, 2018, a single wild salmonberry plant from the backyard of a research team member in Redmond WA was sourced for leaf tissue. RNA was extracted from this leaf tissue and used for transcriptome sequencing and subsequent analysis.

Total RNA from salmonberry leaves was isolated based on a modified protocol of the Plant/Fungi Total RNA Purification Kit (Cat No. 25800) from Norgen Biotek (Ontario, Canada). At harvest, fresh plant leaves were immediately flash frozen in liquid nitrogen, distributed into pre-chilled 50 mL falcon tubes and stored at −80℃ until use. On the day of RNA extraction, the frozen leaf sample was quickly transferred to a pre-chilled mortar, covered in liquid nitrogen and pulverized to a fine powder using a pre-chilled pestle. 100 ug of the powder was transferred to pre-chilled Eppendorf tubes (50 ug in each tube), and 600 uL of lysis buffer was added to each tube. The lysate tube was then incubated at 55℃ for 5 mins with intermittent inverting of the tube. The next steps included clearing of the lysate via column filtration, addition of 100% ethanol (EtOH) to the flowthrough, which was then passed through the grey-ring column for binding. To remove genomic DNA contamination, an on-column RNAse-free DNAse (Qiagen Cat no. 79254) digestion was performed for 15 mins at room temperature. This was followed by column wash steps (three times) and then final elution of total RNA in 50 uL of EB buffer as outlined in the manufacturer’s protocol. In order to obtain a sequencing-ready RNA sample, the total RNA isolate was further subjected to a clean-up protocol developed as described below.

The total RNA isolate (50 uL) in TE buffer was reconstituted in nuclease-free water (50 uL). To this a solution, 600 uL of a 1:1 mix of 100% EtOH and Lysis buffer C (from the Norgen kit) was added. This entire RNA containing mix was then passed through the grey O-ring column via centrifugation. The flowthrough was discarded. The column was next washed three times following the manufacturer’s protocol of the kit. Final elution of the total RNA was then completed in 50 uL of EB buffer. This eluted sample of total RNA was then analyzed by Nanodrop and gel electrophoresis prior to RNA-seq.

### RNA-sequencing and Transcript Assembly

The first round of RNA-seq was conducted by Covance Genomics(Redmond, WA facility). RNA integrity was determined using the 2100 BioAnalyzer (Agilent Technologies, CA) and aRNA Integrity Number (RIN) above 6 was used to generate Illumina sequencing libraries.

The second round of RNA-seq was done by Novogene Corporation Inc. at their Davis facility. The sequencing was performed using Illumina NovaSeq platform (HWI-ST1276) with paired-end 150bp sequencing strategy. Three step quality control (QC) was performed using (1) Qubit 2.0: tests the library concentration preliminarily, (2) Agilent 2100: tests the insert size, (3) Q-PCR: quantifies the library effective concentration precisely. After QC, mRNA is enriched using oligo(dT) beads. First, the mRNA is fragmented randomly by adding fragmentation buffer, then the cDNA is synthesized by using mRNA template and random hexamers primer, after which a custom second-strand synthesis buffer (Illumina), dNTPs, RNase H and DNA polymerase I are added to initiate the second-strand synthesis. Second, after a series of terminal repair, A ligation and sequencing adaptor ligation, the double-stranded cDNA library is completed through size selection and PCR enrichment.

Sequencing raw data for this paper (Table 1) is available from the NCBI SRA database (Bioproject PRJNA1434658, https://www.ncbi.nlm.nih.gov/sra/PRJNA1434658).

**Table 1.**
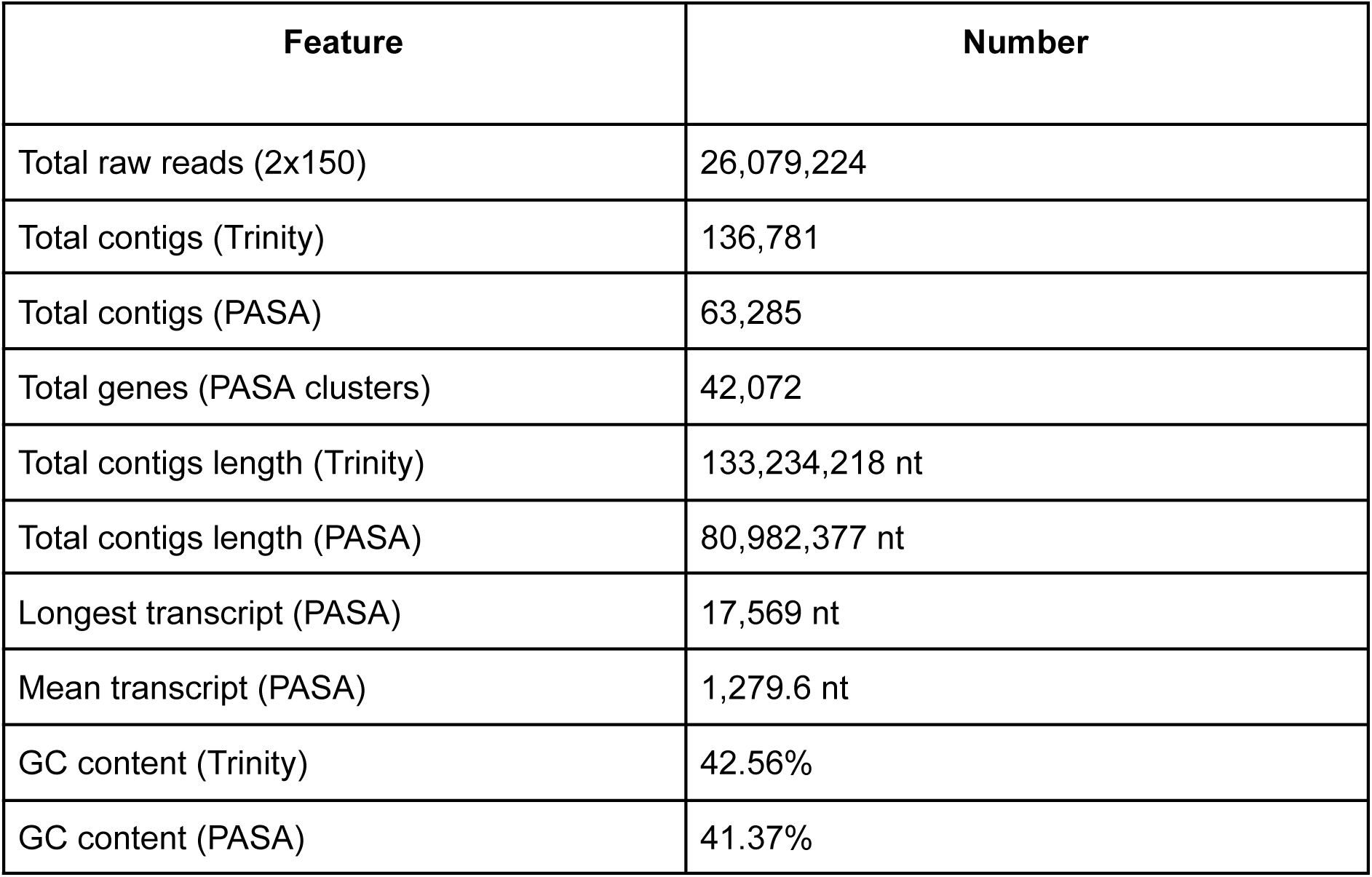
Summary of transcriptome. Sequencing raw reads are deposited at the NCBI SRA database (Bioproject PRJNA1434658, https://www.ncbi.nlm.nih.gov/sra/PRJNA1434658).

| Feature | Number |
| --- | --- |
| Total raw reads (2x150) | 26,079,224 |
| Total contigs (Trinity) | 136,781 |
| Total contigs (PASA) | 63,285 |
| Total genes (PASA clusters) | 42,072 |
| Total contigs length (Trinity) | 133,234,218 nt |
| Total contigs length (PASA) | 80,982,377 nt |
| Longest transcript (PASA) | 17,569 nt |
| Mean transcript (PASA) | 1,279.6 nt |
| GC content (Trinity) | 42.56% |
| GC content (PASA) | 41.37% |

RNAseq data from both rounds of sequencing were assembled together into a set of transcripts in a single FASTA file with the Trinity assembly program (Grabherr et al., 2011) using default settings. The assembly produced 136,781 sequences with a total length of 133,234,218 nucleotides (nt).

### Refining De Novo Transcripts with PASA

Using the Program to Assemble Spliced Alignments (PASA) pipeline (v2.5.3) (Haas, 2008), Trinity-assembled transcripts were filtered and mapped to the salmonberry reference genome (GCA_046118765.1) published by the Canadian BioGenome Project (Sayers et al., 2024). Alignments were performed with Minimap2 (v2.30-r1287) (H. Li, 2021) and pblat (v36×2), (M. Wang & Kong, 2019) and SQLite (v2.8.17) was used to store alignment results. All tools were run with default parameters.

PASA was used to identify and remove redundant and spurious transcripts. The 136,781 Trinity transcripts were reduced to 63,285 transcripts following PASA processing. This filtering step served two primary purposes: minimizing potential contamination from non-salmonberry sequences and merging redundant transcript assemblies generated by Trinity.

### Identifying Putative lncRNA Transcripts

To identify putative long noncoding RNA (lncRNA) transcripts, we annotated the reference salmonberry genome from the Canadian BioGenome Project using Helixer with the *land_plant_v0.3_a_0080.h5* model (Holst et al., 2025). PASA-derived transcripts were mapped to the genome using GMAP (v2021-12-27) run with pruning level 3 (T. D. Wu & Watanabe, 2005).

Putative lncRNA transcripts were initially filtered using FEELnc_filter.pl (v0.2.1) with default settings (Wucher et al., 2017). Coding potential was then evaluated using three independent tools. First, CPAT-plant (v1.2.4) was applied with the Plant-Lnc-pipeline-v2 hexamer model and a coding probability threshold of 0.46 (L. Wang et al., 2013) (Tian et al., 2025). Second, LncFinder (v1.1.6) (Han et al., 2018) was run using training data provided by the Plant-LncRNA-pipeline GitHub repository. Third, PlantLNCBoost (Tian et al., 2025) was run with a classification threshold of 0.5.

To further exclude protein-coding sequences, the filtered candidate lncRNAs were queried against the entire SWIS-PROT protein database (Bateman et al., 2024) (downloaded on June 16, 2025) using DIAMOND (v2.1.11) (Buchfink et al., 2021) with default parameters. Using this information, the PlantLncRNA v2 pipeline script was used to identify a set of high-confidence lncRNA transcripts. Subsequently, putative lncRNA transcripts were analyzed with FEELnc_classifier using the Helixer generated annotations.

### Building Nuclear Phylogenetic Tree

Orthofinder (3.1.2) (Emms & Kelly, 2015) was run on a total of 24 CDS sets from annotated genomes of the Rosaceae family (8 from *Rubus*, 4 from *Rosa*, 12 from *Fragaria)* in addition to salmonberry PASA transcripts. All annotated genomes were obtained from NCBI or rosaceae.org. Diamond (2.1.10) (Buchfink et al., 2021) was used to detect orthologues, Famsa (2.4.1-45c9b26) (Gudyś et al., 2025) was used for alignment, and Fasttree (2.2.0) (Price et al., 2010) for treebuilding. Based on 1,344 trees built for the detected orthogroups, STAG (v1.0.0) (Emms & Kelly, 2018) generated a phylogenetic tree which was displayed with the ETE-3 toolkit’s online web interface (Huerta-Cepas et al., 2016).

### Building Plastome-based Phylogenetic Tree

BLAST homology searches against the *Rubus spectabilis* reference genome identified scaffold JBHWAF010000012.1 as the plastid genome sequence. This plastome was aligned with plastid genomes from 61 *Rubus* species using a circular genome alignment tool, and the resulting alignment was analyzed with PGR-TK (Chin et al., 2023). PGR-TK performed all-versus-all sequence comparisons to identify conserved genomic segments shared among the plastid genomes and generated a cladogram based on shared plastid sequence conservation.

### Identifying Similarities to Rubus Yellow Net Virus Genome

To identify potential viral sequences associated with the salmonberry transcriptome, unmapped Trinity-assembled transcripts were screened using an automated pipeline that performed BLASTN searches against the NCBI nucleotide database. Multiple transcripts showed significant similarity to Rubus yellow net virus (RYNV). To determine whether additional *Rubus*-infecting viruses were present, all publicly available *Rubus* virus genomes were downloaded from NCBI (May 2025) and queried against the complete transcriptome using BLASTN with default parameters; no significant matches other than RYNV were detected.

Protein sequences corresponding to annotated RYNV open reading frames (ORFs) were downloaded and queried against the RYNV-like transcripts using TBLASTN. Transcript length, alignment score, and ORF coverage were used to identify the most complete representative viral sequence, and ORF alignments were compared with the reference RYNV genome (KF241951).

To identify endogenous viral elements in the *Rubus spectabilis* reference genome, RYNV protein sequences were queried against the genome using TBLASTN. Regions corresponding to ORF1 and ORF3 on chromosomes 1 and 3 were extracted with SAMtools faidx (v1.21) (Danecek et al., 2021) and translated using EMBOSS sixpack (v6.6.0.0) (Hancock & Bishop, 2004). Protein reading frames were identified using CD-HIT (v4.8.1) (Fu et al., 2012) with a 40% identity threshold against known badnavirus proteins.

For phylogenetic analysis of ORF1 and ORF3, multiple sequence alignments were generated with MAFFT (v7.526) (Katoh et al., 2017) using the global pair option and 1000 iterations, trimmed with trimAl (v1.5.rev1) (Capella-Gutiérrez et al., 2009) using the automated1 setting, and used to construct phylogenetic trees with IQ-TREE (v3.0.1) (Wong et al., 2025).

## Results

### De novo assembly and annotation of *R. spectabilis* transcriptome from leaves

After sample collection and sequencing, there were a total of 26,079,224 paired-end raw reads. These reads were assembled with Trinity, producing 136,781 contigs, 90,387 (66%) of which were aligned to the salmonberry reference genome with PASA. The Trinity assembly was refined with PASA using the salmonberry genome as a reference, leaving 63,285 contigs, and an estimated 42,072 PASA clusters (Table 1). Because the reference genome contained contigs for non-nuclear organelles, 194 PASA transcripts and estimated genes corresponding to extra-nuclear sequences were excluded from further analysis.

Among the estimated 61,896 non-lncRNA transcripts, 51,330 (83%) were conserved across all 11 publicly available *Rubus* genomes (Table S1), 9,338 were conserved in at least one non-salmonberry *Rubus* genome, and 1,228 transcripts had no match to any other *Rubus* genome.

Eighty-eight (88) sequences were conserved between salmonberry and plants in the *Malus*, *Fragaria*, and *Rosa* genera, but not conserved in any other *Rubus* plant genome besides *R. chamaemorus*. Fifteen (15) nonconserved sequences are longer than 1,000 nt, and their closest matches are described in Table 2. Three (3) of these long sequences had over 90% nucleotide identity matches to *Rosa rugosa*, *Rosa chinensis*, and *Fragaria vesca* (Table 2).

**Table 2.** List of PASA sequences with no homology to other *Rubus* sequences (besides *chamaemorus*.). The three closest matching species are listed in order of highest total bitscore alignment as reported by online blast queries, *Rubus chamaemorus* is excluded. Notes column includes additional information.

| PASA ID | LEN(bp) | First species | Second species | Third species | Notes |
| --- | --- | --- | --- | --- | --- |
| 57059 | 2196 | *Prunus itosakura* | *Potentilla anglica* | *Prunus jamasakura* |  |
| 55771 | 1520 | *Rosa chinensis* | *Rosa rugosa* | *Fragaria vesca* |  |
| 54485 | 2114 | *Rosa rugosa* | *Rosa chinensis* | *Fragaria vesca* | Highly conserved, >90% identity |
| 57059 | 2196 | *Prunus itosakura* | *Potentilla anglica* | *Prunus jamasakura* |  |
| 47665 | 1869 | *Rosa rugosa* | *Rosa chinensis* | *Argentina anserina* | No dna, >90% identity |
| 45791 | 1009 | *Rosa rugosa* | *Rosa chinensis* | *Fragaria vesca* | All >89% identity |
| 44268 | 1728 | *Rosa rugosa* | *Rosa chinensis* | *Fragaria vesca* | >90% identity |
| 43624 | 1058 | *Rosa chinensis* | *Rosa rugosa* | *Fragaria vesca* |  |
| 42445 | 2094 | *Prunus itosakura* | *Potentilla anglica* | *Rosa rugosa* | distant |
| 35747 | 1851 | *Rosa chinensis* | *Fragaria vesca* | *Argentina anserina* | less distant |
| 35545 | 3116 | *Malus domestica* | *Rosa chinensis* | *Malus sylvestrus* |  |
| 35544 |  | *Cotoneaster nebrodensis* | *Malus domestica* | *Malus sylvestris* |  |
| 25369 | 2137 | *Potentilla anglica* | *Rosa rugosa* | *Rosa chinensis* | distant |
| 19711 | 1348 | *Rosa chinensis* | *Rosa rugosa* | *Potentilla anglica* | distant |
| 17478 | 1089 | *Rosa rugosa* | *Rosa chinensis* | *Fragaria vesca* | No dna, >90% identity |
| 1553 | 2237 | *Potentilla anglica* | *Rosa chinensis* | *Fragaria vesca* |  |

### Detection of lncRNA sequences and analysis of putative lncRNA transcripts

Long non-coding RNAs (lncRNAs) are RNA sequences that are at least 200 nts long, and do not code for any proteins. In plants, lncRNAs often have critical roles throughout development and in response to environmental cues, such as controlling occupation rates of gene promoter regions, chromosome looping, chromatin modification, and miRNA biogenesis (Wierzbicki et al., 2021). Due to the inherent difficulty in determining whether a particular lncRNA sequence is functional, interspecies conservation is often used as a proxy for functionality, and lncRNAs are significantly less conserved than mRNA sequences (Deng et al., 2017).

Out of the 63,285 assembled PASA transcripts from salmonberry, 62,294 (98.4%) had length greater than or equal to 200 nts, of these, 1,695 (2.7%) were identified as potential lncRNAs by FEELnc_filter. A comparison of three different lncRNA identification tools/software identified a consensus of lncRNAs that did not encode known proteins based on SWIS-PROT. This analysis identified 1,389 high-confidence putative lncRNA transcripts (Fig. 1).

**Fig 1.**
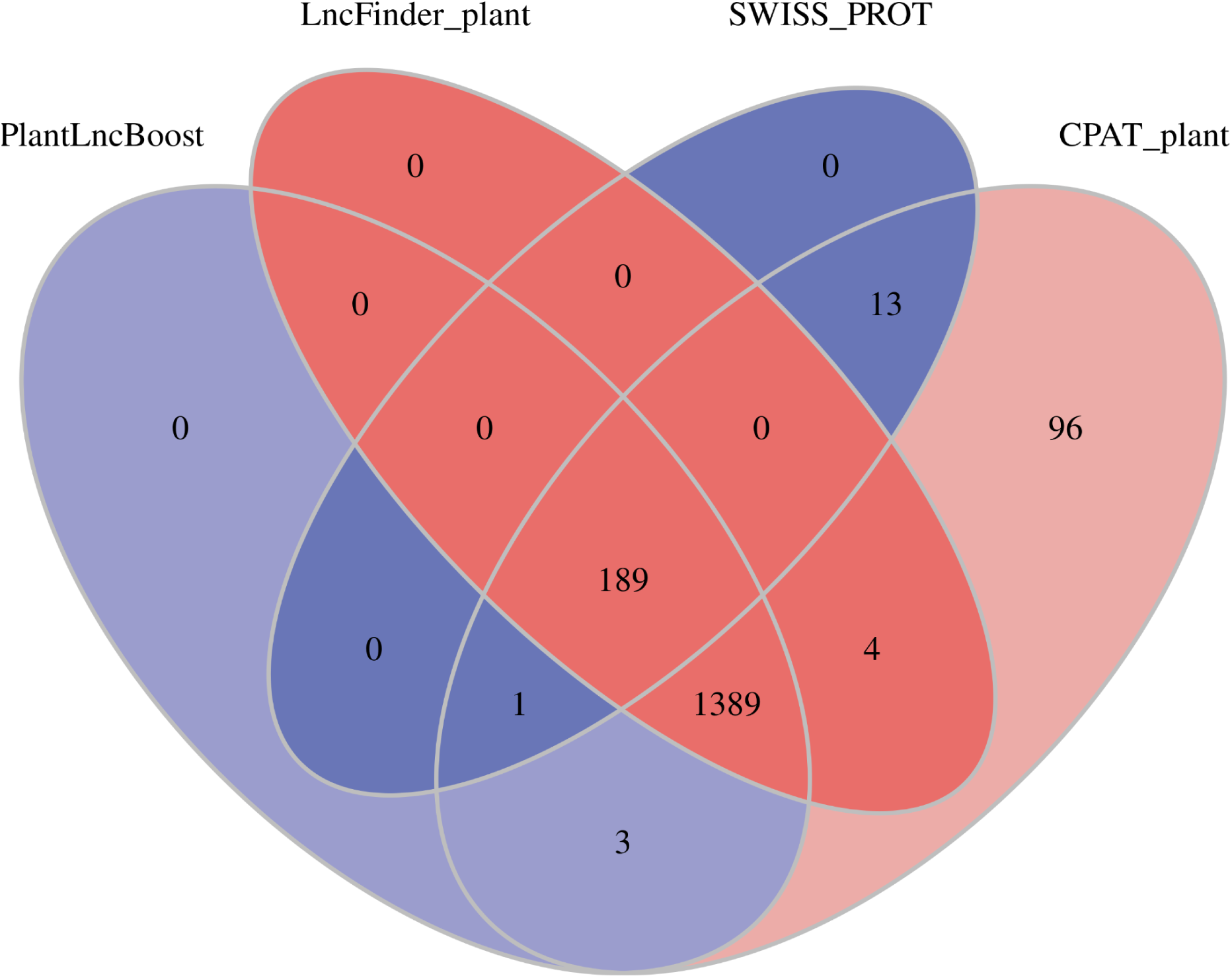
Venn diagram of various tools identifying lncRNA transcripts. Transcripts that have been identified as lncRNAs by all three tools, without matching any known proteins, are considered high-confidence putative lncRNA transcripts.

After finding 1,389 high-confidence putative lncRNA transcripts, FEELnc_classifier categorized these transcripts with respect to the Helixer annotated genome (Holst et al., 2025). Herein, we use FEELnc’s definitions for the various categories. For example, “intergenic” lnRNA transcripts are transcripts that do not overlap a predicted protein-coding gene, whereas “nested” lncRNA transcripts are contained within a predicted protein-coding gene, often as an intron (Wucher et al., 2017). Based on the Helixer annotation, 1127 (81%) putative lncRNA transcripts expressed in leaves of salmonberry are intergenic (Fig. S1). This is consistent with previously reported observations, where over 50% of plant lncRNAs are intergenic (Bedre, 2024). Of these 1,389 high-confidence putative lncRNA transcripts, 218 transcripts are heavily conserved between all 11 sequenced *Rubus* genomes, 731 transcripts were conserved in at least one other *Rubus* genome and the remaining 440 not conserved in any other *Rubus* genome.

## Relation to Other Rosaecea and *Rubus* Species

### Phylogenetic Analysis of Conserved Nuclear Sequences

A total of 24 genomes and their associated annotations were downloaded from the NCBI database and the Genome Database for Rosaceae. Orthofinder3 was run against the coding sequences extracted from these 24 annotated genomes, and against the PASA transcripts. Fig. 2 shows the resulting phylogenetic tree produced by Orthofinder3 using STAG (Species Tree Inference from All Genes,) based on 1,344 orthogroups conserved across all 25 genomes and transcriptomes. Among the *Rubus* species with a complete, annotated genome, salmonberry is most closely related to *Rubus ellipticus* and *Rubus chingii hu*.

**Fig 2.**
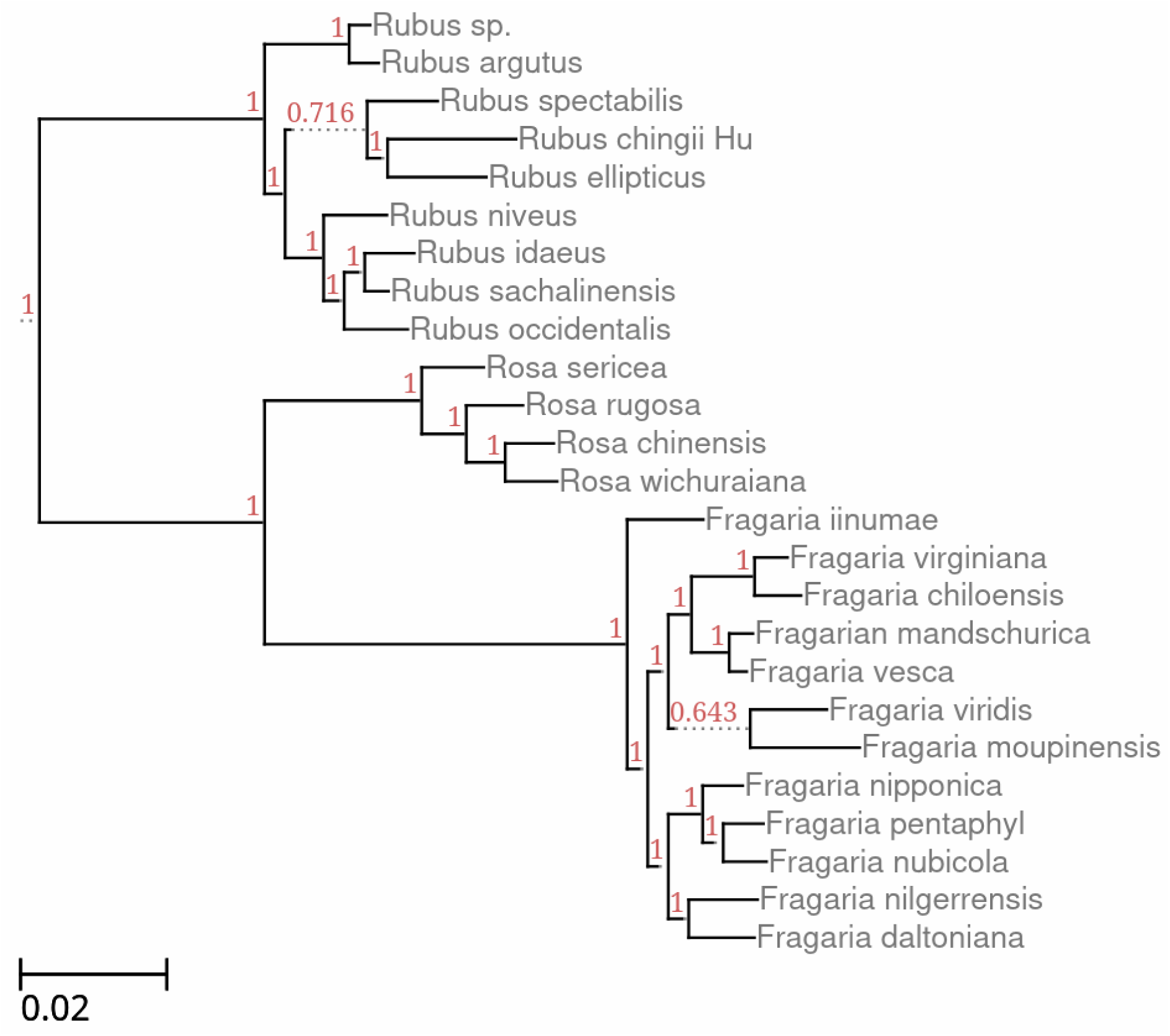
Orthofinder consensus tree. Consensus tree of plant species from *Rubus*, *Rosa*, and *Fragaria* genera based on annotated protein-coding genes built using Orthofinder.

### Phylogenetic Analysis of *Rubus* Plastomes

Plastid genome sequences for *Rubus* species were downloaded from the National Center for Biotechnology Information (NCBI) database and combined into a single FASTA file, to which the salmonberry plastid genome was added. This dataset was analyzed using PGR-TK (Chin et al., 2023) to assess plastid genome conservation and structural variation.

Among all surveyed species, salmonberry is closest to *Rubus arcticus*. This confirms previously reported findings by Carter, 2019, where *R. Arcticus, R. Spectabilis, R. Hawainsis,* and *R. Pubescens* were found to be a monophyletic subgroup of *Rubus* using both plastome and nucleolar based methods. It is impossible to properly compare Fig. 2 and Fig. 3 because there are few fully annotated *Rubus* genomes relative to the number of sequenced plastomes.

**Fig 3.**
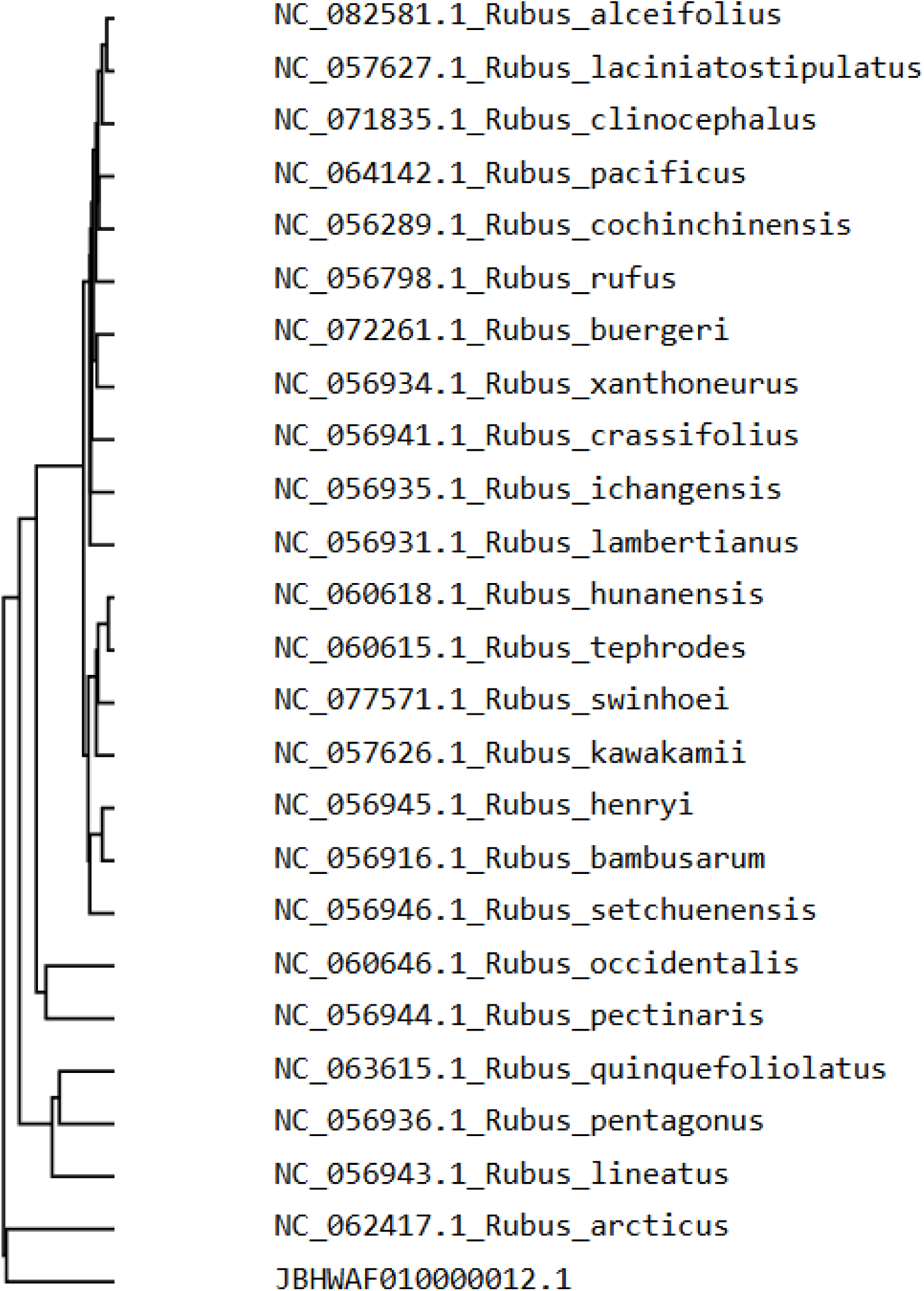
PGR-TK phylogeny tree. The phylogeny tree of plant species from *Rubus* is built from plastid sequences using PGR-TK. JBHWAF010000012.1 represents salmonberry (*Rubus spectabilis*).

### Identification of a New Endogenous Virus Species in Salmonberry

Rubus yellow net virus (RYNV) is a member of the family Caulimoviridae and infects species within the genus *Rubus*, including cultivated raspberries and blackberries (Jones et al., 2002) (Vakic et al., 2022). RYNV infections may cause foliar symptoms such as yellow vein banding or net-like chlorosis. Importantly, RYNV can exist as episomal viral particles or as endogenous viral sequences integrated into host genomes, complicating molecular detection and interpretation (Ho et al., 2024).

Analysis of the salmonberry transcriptome revealed seven (7) Trinity transcripts (200–6,500 nt in length) with significant nucleotide similarity to raspberry RYNV. Among these, transcript TRINITY_DN6392_c0_g1_i1 was the longest (6,482 nt), showed the highest-scoring BLASTN match to the RYNV genome, and exhibited the strongest protein-level similarity to RYNV ORFs.

TBLASTN analysis demonstrated that six of the seven RYNV protein-coding ORFs (ORFs 1 and 3–7) aligned significantly to TRINITY_DN6392_c0_g1_i1 (Table 3). Notably, no similarity was detected to ORF2, an ORF that is conserved across all known badnaviruses and is commonly used as a marker for episomal infections (Bhat et al., 2023) (Selvarajan et al., 2016). In addition, the length of TRINITY_DN6392_c0_g1_i1 is shorter than the full RYNV genome (7,932 bp) and shorter than the smallest known badnavirus genome (Cacao swollen shoot virus, 7,006 bp) (Bhat et al., 2016). Together, these observations indicate that TRINITY_DN6392_c0_g1_i1 represents an incomplete viral genome fragment rather than a complete episomal virus.

**Table 3.** ORF locations in badnavirus. Comparison of ORF start and end locations in the Salmonberry Trinity transcript, and their position in the RYNV genome.

| ORF | RYNV protein start (nt) | RYNV protein end (nt) | Genome strand | Transcript strt (nt) | Transcript end (nt) | Transcript Strand |
| --- | --- | --- | --- | --- | --- | --- |
| ORF1 | 386 | 1018 | + | 7 | 579 | - |
| ORF2 | 1015 | 1476 | + | N/A | N/A | N/A |
| ORF3 | 1433 | 7348 | + | 1512 | 6455 | - |
| ORF4 | 6940 | 7356 | + | 1576 | 1782 | - |
| ORF5 | 7654 | 180 | + | 4290 | 4556 | + |
| ORF6 | 3330 | 3767 | - | 4290 | 4715 | + |
| ORF7 | 7906 | 405 | - | 4164 | 4556 | - |

Inspection of the Canadian BioGenome Project salmonberry reference genome revealed multiple endogenous RYNV-like integrations. Two adjacent copies of a full-length RYNV-like genome were identified on chromosome 3. In both copies, ORF1 shared 87–88% nucleotide identity with TRINITY_DN6392_c0_g1_i1, while ORF3 shared 82% nucleotide identity.

Additionally, a distinct but closely related endogenous badnavirus species was identified on chromosome 1, represented by two genomic copies. These copies exhibited 78% nucleotide identity in ORF1 and 77% identity in ORF3 relative to the Trinity transcript, falling below the 80% nucleotide identity threshold commonly used to define viral species boundaries and indicating a distinct viral species. However, both integrated lineages have over 80% ORF1/ORF3 nucleotide identity with each other. Fig. 4 shows a phylogenetic tree of all novel viral lineages compared with already-known badnaviruses using ORF3 protein alignment, Fig. S2 is a phylogenetic tree based on ORF1.

**Fig 4.**
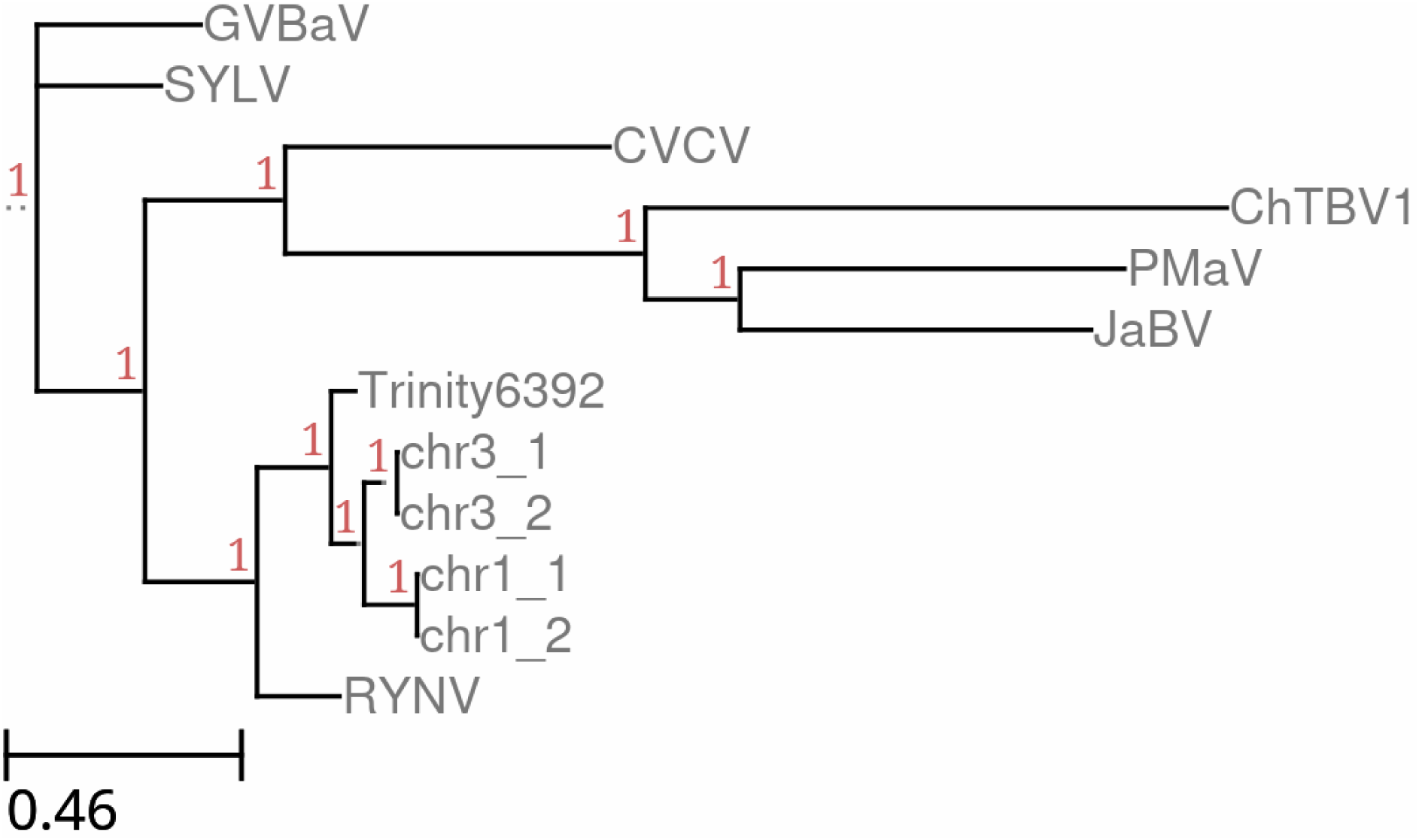
Phylogenetic tree based on ORF3 for a variety of the known badnaviruses. GVBaV (Gooseberry Vein Banding associated Virus,) SYLV (Spiraea Yellow Leafspot Virus,) CVCV (Codonopsis Vein Clearing Virus) ChTBV1 (Chinaberry Tree Badnavirus 1,) PMaV (Pandanus Mosaic associated Virus,) RYNV (Rubus Yellow Net Virus,) and JaBV (Jujube-associated Badnavirus.) Trinity6392 is the viral sequence found in the Trinity assembly, chr3_1 is the first viral sequence in the third chromosome of the salmonberry reference genome, with a similar naming scheme for chr3_2, chr1_1, chr1_2.

To assess the broader distribution of these viral sequences, BLAST searches were performed against all 11 publicly available *Rubus* nuclear genome assemblies. Low scoring matches to the salmonberry endogenous viral regions were detected in *R. idaeus* and *R. sachalinensis*, but these integrations occurred on different chromosomes and genomic locations, suggesting independent integration events rather than inheritance from a common ancestor.

## Discussion

This study presents the first comprehensive leaf transcriptome analysis of *Rubus spectabilis*, integrating transcriptomic, comparative genomic, plastid phylogenetic, and viral discovery approaches to better resolve its molecular biology and evolutionary context within *Rubus*.

### Transcriptome Preparation

The RNeasy Plant Mini Kit yielded RNA of insufficient quality for sequencing, whereas the Plant/Fungi Total RNA Purification Kit (Norgen Biotek) produced high-quality RNA suitable for library preparation. This likely reflects more effective removal of secondary metabolites and other inhibitors, highlighting the importance of protocol optimization for metabolite-rich woody taxa within the Rosaceae.

Transcriptome refinement using PASA substantially reduced redundancy and possible contaminants, collapsing 136,781 Trinity contigs to 63,285 transcripts grouped into 42,072 clusters. While this improved assembly coherence, it may also have led to the loss of biologically meaningful variation. Notably, the reference genome derives from a geographically distinct individual (Vancouver, Canada), whereas our sample originated from the Seattle region. Such geographic divergence may contribute to reduced mapping efficiency and the exclusion of unconserved or population-specific transcripts during refinement.

Consistent with this, only 42,697 transcripts matched coding sequences predicted by Helixer and LiftOff (based on *Rubus chingii hu* annotations), leaving 20,588 mapped but unannotated transcripts. While this limitation does not substantially affect phylogenetic inference or viral detection, it reduces confidence in lncRNA characterization and highlights the need for improved genome annotation in salmonberry.

### Phylogeny

We reconstructed phylogenetic relationships using both plastome and nuclear datasets. Plastome-based analysis, generated from a broad sampling of *Rubus* plastid genomes, likely provides a more robust estimate of species relationships due to its completeness and reduced sensitivity to gene duplication. In contrast, nuclear phylogenetic reconstruction using Orthofinder was constrained by limited availability of fully annotated genomes across Rosaceae taxa.

Additional challenges arose from the use of transcript-derived sequences, where alternative splicing can introduce artificial gaps, and from the high prevalence of gene duplication in Rosaceae. The absence of single-copy orthologs and reliance on multi-copy gene families increase error rates in species tree inference (Emms & Kelly, 2018). These limitations underscore the need for more complete nuclear genome resources across *Rubus* and related genera.

### Viral and symbiotic associations

We identified a putative novel virus associated with salmonberry, phylogenetically related to Rubus yellow net virus. However, its biological status remains unclear. It is unknown whether the detected sequences represent active infection or endogenous viral elements integrated into the host genome. Future work using RT-PCR across open reading frames, small RNA profiling, and viral particle isolation will be necessary to distinguish between these possibilities and to assess potential impacts on host physiology.

In addition to viral sequences, internal transcribed spacer (ITS) data revealed the presence of multiple fungal endophytes, including *Albifimbria verrucaria*, *Septoria rosae*, and *Cladosporium spp*. These taxa include both pathogenic and potentially mutualistic species (Yin et al., 2023; Bagsic et al., 2015; Răut et al., 2021), suggesting a complex leaf microbiome. Systematic characterization of these communities represents an important avenue for future research.

### Implications and future directions

Together, these results expand genomic resources for *Rubus spectabilis* and provide new insight into transcriptome diversity, noncoding RNA conservation, phylogenetic relationships, and host-associated viruses within *Rubus*. The conserved subset of high-confidence lncRNAs identified here likely plays important functional roles across the genus and warrants further investigation. Similarly, transcripts unique to salmonberry yet conserved in more distantly related taxa may represent lineage-specific innovations with potential relevance for crop improvement in cultivated *Rubus* species.

Future efforts should prioritize improved genome annotation, population-level sampling to capture intraspecific variation, and functional validation of candidate genes and noncoding RNAs. Collectively, such work will deepen our understanding of evolutionary and functional genomics within this ecologically and economically important group.

## Acknowledgements

We gratefully acknowledge Kerry Deutsch and LabCorp Inc. for performing RNA-seq on our samples and B. Platt for his assistance in the project. We also thank Lisa and John Platt for providing meeting space to the Salmonberry Research Group, and Dan McReavy, Lisa/John Platt, Dharmesh Syal, Robin Carmichael, Chris Amemiya, Petros Bournelis, Jay Rajagopalan, and Jennifer Salley for their generous support of the project through IndieGoGo or directly.

## Author Contributions

MPS, MB and JG conceived the project. All authors contributed to organizing the crowdfunding effort. MB and TM planned the experiments. All authors conducted the experiments. ALS, NN, AR and MPS performed the analysis. ALS, JG, MB and MPS wrote the paper. JG managed the project.

## Data Availability

Raw paired-end reads are available from the Short Read Archive hosted by NCBI under Bioproject PRJNA1434658, or at the URL https://www.ncbi.nlm.nih.gov/sra/PRJNA1434658. The de-novo Trinity and refined PASA assemblies are available as supplementary files under rSpecTrinity.fa and rSpecPasa.fa. The gff3 annotation files generated by PASA and Helixer are available as supplementary files rSpecPasa.gff and rSpecHelixer.gff. The text file listing all high-confidence lncRNA PASA transcript IDs is saved as Final_lncRNA_results.txt and the corresponding file generated by FEELnc_classifier is stored as lncRNA_classes.csv. Finally, the file containing all of the PASA transcript ids with no detected homology to any other Rubus genome (besides R. chamaemorus) is stored in rSpecNoRubusHomology.txt.

## Notes

https://www.ncbi.nlm.nih.gov/sra/PRJNA1434658

### Competing Interest Statement

The authors have declared no competing interest.

