## supplemental tables and figures for "Salmonberry Transcriptome Reveals Phylogeny and Novel Badnavirus Species"

### Supplementary Material

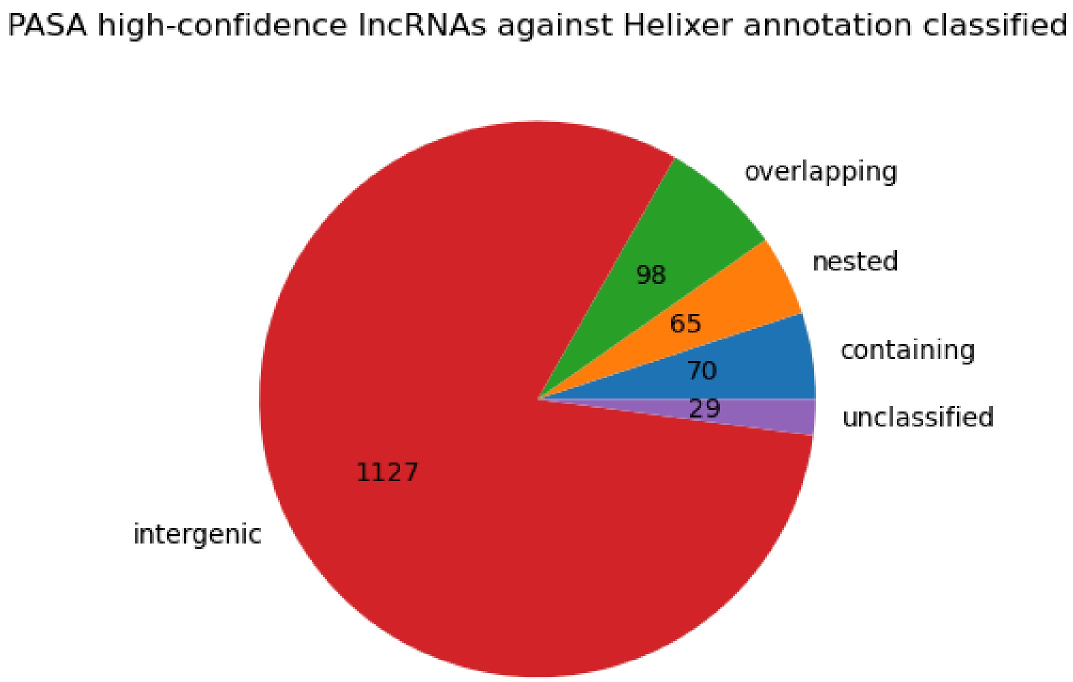


**Fig. S1. Visualized output from FEELnc_classifier ran on high-confidence putative lncRNA transcripts**. All categories are mutually exclusive.

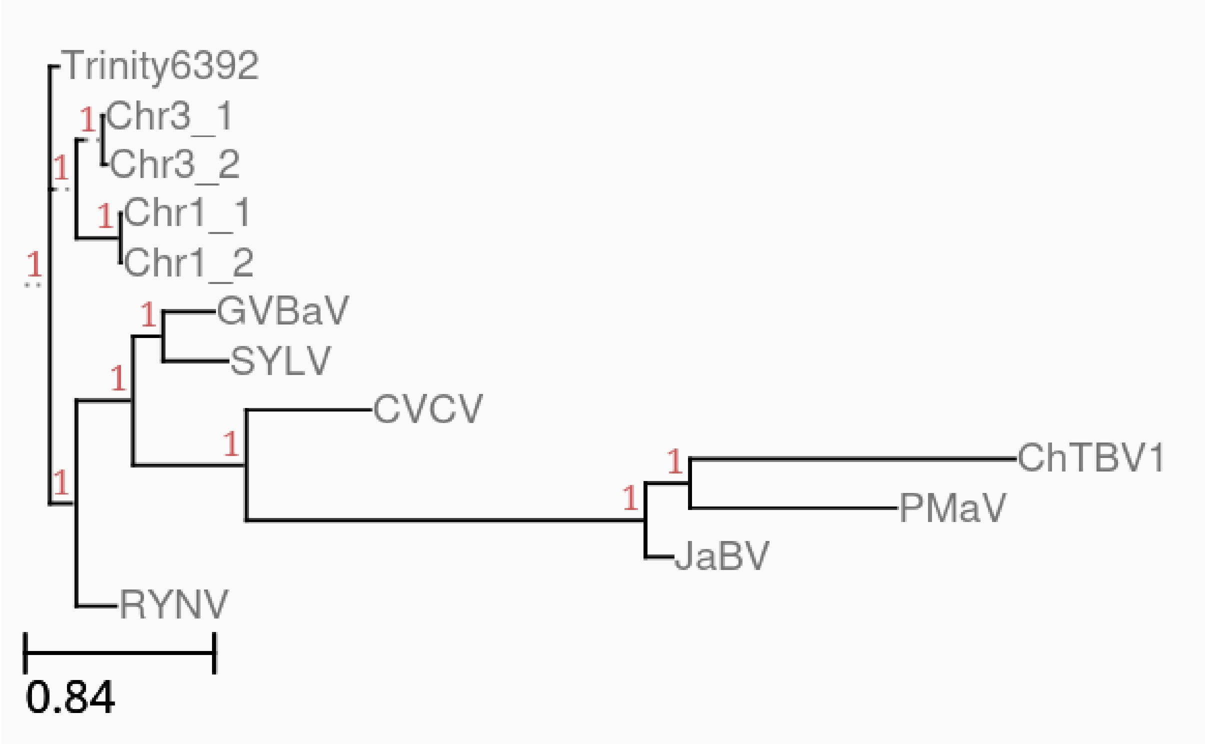


**Fig. S2. Phylogenetic tree based on ORF1 for a variety of the known badnaviruses** GVBaV (Gooseberry Vein Banding associated Virus,) SYLV (Spiraea Yellow Leafspot Virus,) CVCV (Codonopsis Vein Clearing Virus) ChTBV1 (Chinaberry Tree Badnavirus 1,) PMaV (Pandanus Mosaic associated Virus,) RYNV (Rubus Yellow Net Virus,) and JaBV (Jujube-associated Badnavirus.) Trinity6392 is the viral sequence found in the Trinity assembly, chr3_1 is the first viral sequence in the third chromosome of the salmonberry reference genome, with a similar naming scheme for chr3_2, chr1_1, chr1_2

| Species | GCA identifier | Year |
|---|---|---|
| *parviflorus* | GCA_029955375.1 | 2023 |
| *caesius* | GCA_964235055.1 | 2024 |
| *niveus* | N/A | 2025 |
| *sachalinensis* | N/A | 2025 |
| *chingii Hu* | N/A | 2021 |
| *argutus* | GCA_040183295.1 | 2024 |
| *ellipticus* | N/A | 2025 |
| *hochstetterorum* | GCA_040496155.1 | 2024 |

| | | |
|---|---|---|
| *alceifolus* | GCA_051990025.1 | 2025 |
| *idaeus* | N/A (Joan J v2) | 2022 |
| *occidentalis* | N/A (v3) | 2018 |

**Table S1. Rubus genome assemblies**. All assemblies without an associated GCA identifier were obtained from rosaceae.org
